# Secondary bile acid lithocholic acid ameliorates colitis-like inflammation in a human intestine-on-chip system

**DOI:** 10.1101/2025.11.07.686950

**Authors:** Tim Kaden, Manuel Allwang, Johannes Stallhofer, Katja Graf, Martin Raasch, Alexander S. Mosig

## Abstract

Inflammatory bowel disease (IBD) is a multifactorial disease of the gastrointestinal tract without curative treatment. Previous studies highlighted that altered fecal bile acid levels correlate with intestinal microbiota composition changes and inflammation in IBD. Lithocholic acid (LCA) is a secondary bile acid (SBA) drastically reduced during active IBD but mediates beneficial effects at the mucosal intestinal barrier during intestinal homeostasis. In a dextran sodium sulfate (DSS)-induced colitis-on-chip model, it was investigated whether the administration of LCA has a protective impact on inflammation-mediated tissue damage. Physiological responses were successfully recapitulated in the human colitis model, enabling the dissection of individual cell responses. Treatment with LCA concentrations similar to healthy human intestinal levels efficiently ameliorated the colitis-like phenotype. LCA treatment stimulated epithelial cell proliferation, thereby maintaining villus morphology, intestinal barrier integrity, and reducing inflammation. The protective effects of LCA were mainly mediated by the activation of the farnesoid X receptor (FXR).

## 1. Introduction

Inflammatory bowel disease (IBD) comprises two main types of chronic diseases of the gastrointestinal tract, namely Crohn’s disease (CD) and ulcerative colitis (UC), which show drastically increasing prevalences worldwide^1^. Both disease types are associated with key pathophysiological features, including intestinal barrier defects, morphological atrophies, excessive mucosal inflammation of the host, and microbial dysbiosis^2^. Despite advancements in therapeutic intervention strategies, no causative curative treatments exist. IBD significantly impairs the quality of life for patients and is also associated with a substantial risk of developing life-threatening extraintestinal manifestations^3^.

IBD is commonly treated with immunomodulatory drugs or biologics to counteract the intestinal inflammation and mitigate the disease symptoms^4^. However, primary nonresponse or secondary loss of response are major limitations to effectively induce disease remission^5^. Many IBD patients require surgery. About 10% of patients with UC have to undergo colectomy due to refractory flares or the development of colorectal cancer as a result of long-standing inflammation^6,7^. Up to 80% of patients with CD develop complications like fibrotic stenoses or fistulas with abscesses that have to be treated by a surgeon at least once in lifetime^8^. Therefore, the emerging paradigms in treating IBD are shifting towards new strategies to improve the underlying microbial dysbiosis by restoring the microecology and a microbial metabolome pool that preserves the host integrity by promoting barrier function and immunomodulation^2,9^.

Dysbiosis of the intestinal microbiota is frequently observed in IBD patients^10^, leading to reduced levels of microbial-derived metabolites, including secondary bile acids (SBAs)^11–13^, which are central in regulating gut homeostasis. These postbiotics bile acids (BAs) are produced through metabolization by intestinal microbes and have been proposed as potential mediators of epithelial barrier integrity and immune responses^14–16^, as well as regulators of microbiota composition by limiting excessive pathogen growth through antimicrobial activity^15,17^.

In this context, lithocholic acid (LCA) is a SBA that is generated in the intestine from liver-derived primary bile acids (PBAs) such as chenodeoxycholic acid and cholic acid mainly by 7α-dehydroxylase-producing *Clostridium* species^18^. A deficiency in SBA-producing bacteria was reported to be associated with onset and progression of UC, while administration of LCA mitigated intestinal inflammation^19^. These findings suggest a causal relationship between reduced LCA levels and IBD progression.

Results from preclinical *in vitro* epithelial monocultures already suggested a potentially protective effect of LCA by reducing inflammation after chemical stimulation with dextran sodium sulfate (DSS)^20^ or IL-1β^21^ and maintaining expression of apical junctional complexes (AJCs) and barrier integrity^22–24^. Mouse models have further provided fundamental results on the effects of LCA to prevent IBD in a more holistic setting. Conventionally, DSS is used in mice to induce experimental colitis^25,26^. Administration of LCA reduced colitis-associated inflammation, inhibited epithelial apoptosis, and decreased the overall disease severity^20,19,27,28^.

Despite many important findings, epithelial monocultures inherently lack the multicellular complexity of native tissues, particularly the presence of immune cells mediating inflammatory cross-talk, and do not accurately replicate physiological tissue barriers and interfaces found *in vivo*^2^. While animal models provide the evaluation of systemic responses, it remains challenging to dissect individual disease-driving factors in a spatial-temporal manner and to unravel cell-specific pathophysiological responses. Furthermore, the extrapolation of data from animal models remains challenging due to cross-species differences^29,30^, in particular regarding host immune responses. Human microphysiological intestine-on-chip (IoC) models are promising tools to overcome some of the current limitations of static 2D models and animal models^2,31^. IoC models emulate relevant aspects of the intestinal microenvironment by integrating multicellularity complexity, peristalsis-induced villus morphogenesis, enhanced barrier integrity, and differentiation of epithelial cells into distinct subtypes^32^.

In this study, a DSS-induced colitis-on-chip model was utilized to study potentially protective LCA effects in the context of IBD. Clinically relevant IBD markers such as inflammation, barrier dysfunction, AJC alterations, and intestinal cell proliferation were investigated, both at the intestinal epithelial cell barrier and the adjacent vascularity including endothelial cells and immune cells. Treatment with specific BA receptor antagonists showed that LCA-associated effects, indicated by maintaining barrier integrity and cell proliferation are linked to the farnesoid X receptor (FXR).

## 2. Results

### 2.1 Emulation of a DSS-induced colitis-like phenotype in the IoC model

A previously described perfused IoC model^33,34^ was used, which reproduces an *in vivo*-like three-dimensional (3D) intestinal microanatomy. The luminal abundance of gram-negative bacteria was mimicked by administration of bacterial lipopolysaccharide (LPS) in the intestinal channel. In agreement with previous results^33^, LPS itself did not reduce intestinal epithelial barrier integrity or triggered adverse cytokine release (**Supplementary Figure S1A**), demonstrating epithelial immunotolerance to the bacterial endotoxin. However, when LPS was administered in the vascular channel, cytokine levels were significantly increased, replicating conditions of endotoxemia (**Supplementary Figure S1B**)^35^. In IBD, intestinal microbes can invade the mucosal barrier and translocate into the systemic blood circulation due to impaired host defense mechanisms and increased epithelial barrier permeability, resulting in chronic inflammation with potentially life-threatening infections affecting additional organs^36^.

To recapitulate a colitis-like phenotype, 1.5% DSS, a concentration that is comparable to experimental models replicating conditions of IBD^26,37,38^, was perfused in combination with LPS in the intestinal channel for 48 h (**Figure 1A,B**). Light microscopic monitoring revealed morphological atrophies in villus-like structures induced by DSS compared to models without treatment (**Figure 1C**). Quantification of the villus height of the intestinal model showed a significant and time-dependent reduction upon DSS treatment (**Figure 1D**). Further, DSS perfusion induced a significant increase in permeability measured by permeation of fluorescein isothiocyanate (FITC)-dextran beads from the luminal intestinal model side into the vascular compartment (**Figure 1E**). However, the overall viability of the tissue models was not found to be compromised by DSS treatment (**Supplementary Figure S2**).

**Figure 1:**
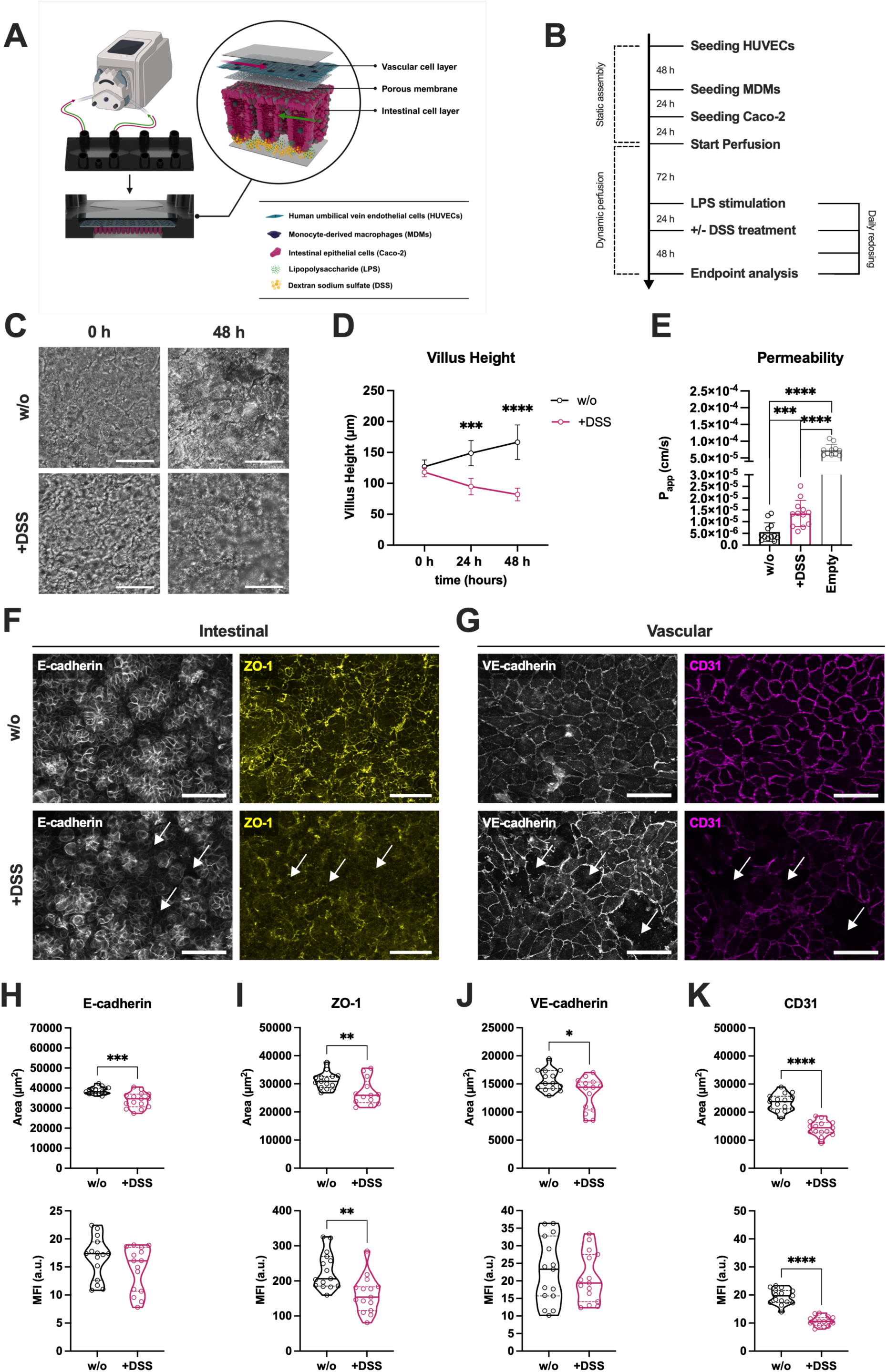
Recapitulating of a colitis-like IBD phenotype in the IoC model. **(A)** Schematic illustration of the colitis-on-chip model. HUVECs (turquoise) and MDMs (indigo) are seeded in the top channel to recapitulate the vascular cell layer. Caco-2 cells (pink) form the intestinal epithelial barrier in the bottom channel of the chip. An integrated porous membrane (grey, with pores) separates both channels. The cellular model depicts an idealized structure of the crypt- and villus-like structures forming in the model after induction of peristaltic perfusion. Both channels were bidirectionally perfused with cell-specific medium. The colitis-like phenotype was triggered by perfusion of DSS (yellow particles) and LPS (green particles) at the luminal side in the bottom intestinal channel. Created with BioRender.com. **(B)** Timeline of static model assembly, perfusion, and experimental manipulation. **(C)** Representative light microscopic images of intestinal epithelial morphology in untreated (w/o) and DSS-treated (+DSS) intestinal models after 0 h and 48 h. Scale bars, 100 µm. **(D)** Monitoring of villus height (in µm) in untreated (w/o) and DSS-treated (+DSS) models over 48 h. Data points represent mean ± SD of 4 independent experiments (n = 4) with 3 individual MDM donors. ***p ≤ 0.001, ****p ≤ 0.0001 (Two-way ANOVA with Šídák’s multiple comparisons test). **(E)** Measurement of FITC-dextran permeability (shown as calculated permeability coefficient P_app_ in cm/s) in untreated (w/o) and DSS-treated (+DSS) models compared to chips without cells (Empty) after 48 h. Bars represent mean ± SD of 12 independent experiments (n = 12) with 7 individual MDM donors. ***p ≤ 0.001, ****p ≤ 0.0001 (Paired two-tailed t test). **(F,G)** Representative immunofluorescence images of intestinal **(F)** and vascular cell layers **(G)** in untreated (w/o) and DSS-treated (+DSS) models after 48 h. Caco-2 cells were stained for E-cadherin (white) and ZO-1 (yellow). HUVECs were stained for VE-cadherin (white) and CD31 (magenta). Scale bars, 100 µm. **(H-K)** Quantification of covered area (upper row, in µm^2^) and mean fluorescence intensities as arbitrary units (lower row, MFI a.u.) of E-cadherin **(H)**, ZO-1 **(I)**, VE-cadherin **(J)**, and CD31 **(K)** signals. Violin plots with indicated median and quartiles of 3 independent experiments (n = 3) with 3 independent MDM donors. 5 different images were taken for each experiment. *p ≤ 0.05, **p ≤ 0.01,***p ≤ 0.001, ****p ≤ 0.0001 (Unpaired two-tailed t test).

Previous studies have demonstrated reduced expression AJCs, such as adherens junction protein epithelial (E)-cadherin and tight junction protein zonula occludens-1 (ZO-1), in UC^39,40^. To recapitulate this clinically relevant disease hallmark, immunofluorescence staining followed by quantification of fluorescence image data using an automated image-analysis pipeline in CellProfiler were performed on samples from untreated and DSS-induced colitis-on-chip models. This allowed to extract and measure fluorescence intensities and the covered signal area of identified AJCs and adhesion molecule networks (**Supplementary Figure S3**). DSS treatment disrupted epithelial and vascular AJCs and adhesion molecules in the IoC model (**Figure 1F,G**), reflected by a significant decrease in the covered area by E-cadherin. Mean fluorescence intensity (MFI) within the segmented E-cadherin network, however, only exhibited a decreasing trend compared to untreated models (**Figure 1H**). Similarly, the covered area and MFI of ZO-1 were diminished in DSS-treated models (**Figure 1I**). Furthermore, a significant reduction of the covered area by adherens protein vascular endothelial (VE)-cadherin was observed after DSS administration (**Figure 1J**). Comparable effects were observed for expression of platelet and endothelial cell adhesion molecule 1 (PECAM-1, CD31) (**Figure 1K**), for which DSS induced a significant decrease in MFI and the area covered by this endothelial adhesion molecule.

3D reconstructions of the tissue further demonstrated that DSS treatment enabled migration of vascularly integrated MDMs through the compromised endothelial barrier and across the porous membrane into the luminal epithelial model side (**Supplementary Figure S4**). In contrast, MDMs in untreated models remained confined to the vascular channel.

### 2.2 Pro-inflammatory cytokine release is elevated upon DSS-induced barrier disruption

To evaluate the inflammatory response in the colitis-on-chip model, the release of pro-inflammatory cytokines IL-6, TNF-α, IL-8, and IL-1β (**Figure 2**), which are typically elevated in colitis patients^41^, was measured. DSS treatment significantly increased TNF-α (**Figure 2B**) and IL-8 (**Figure 2C**) in the vascular compartment, while IL-6 showed an upward trend. In the intestinal compartment, DSS led to a significant rise in TNF-α (**Figure 2F**) and IL-1β (**Figure 2H**), while IL-6 and IL-8 showing non-significant trends toward elevation (**Figure 2E,G**).

**Figure 2:**
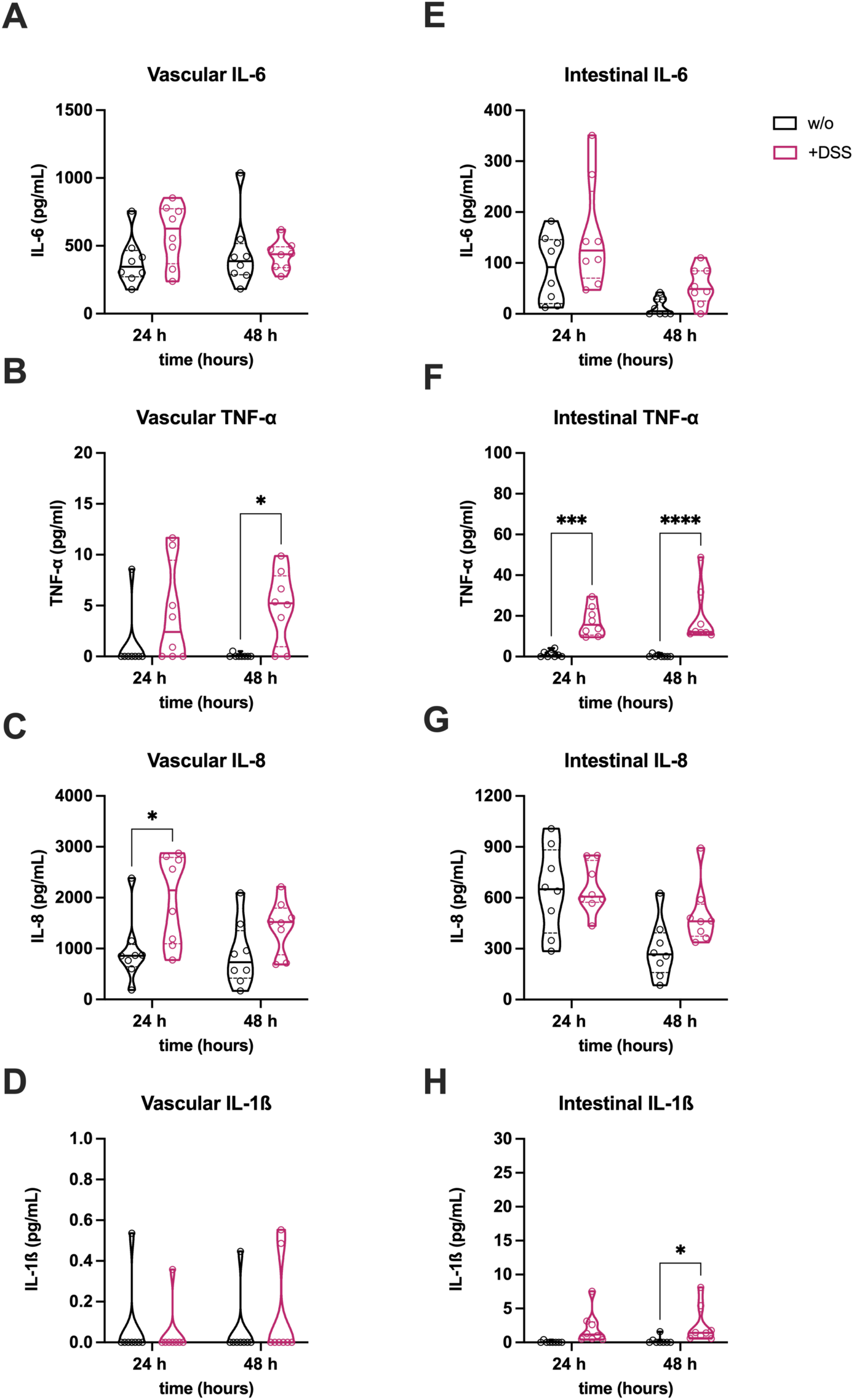
DSS induces inflammation in the IoC model. Release of pro-inflammatory cytokines (IL-6, TNF-α, IL-8, IL-1β) in vascular **(A-D)** and intestinal **(E-H)** medium supernatants of untreated (w/o) and DSS-treated (+DSS) models. Violin plots with indicated median and quartiles of 8 independent experiments (n = 8) with 6 individual MDM donors. *p ≤ 0.05, ***p ≤ 0.001, ****p ≤ 0.0001 (Two-way ANOVA with Šídák’s multiple comparisons test).

### 2.3 LCA alleviates the colitis-like phenotype by preserving intestinal 3D morphology and barrier integrity

Effects of LCA on cell viability were investigated at different concentrations in 2D cell cultures of Caco-2 and HUVECs. It was shown that LCA did not compromise cell viability up to 100 µM in Caco-2 cells (**Supplementary Figure S5A**). In contrast, HUVECs exhibited a significant reduction in viability at LCA concentrations ≥ 50 µM (**Supplementary Figure S5B**). To investigate the potential of LCA in ameliorating inflammation-associated effects in the colitis-on-chip model, it was administered at a human-relevant fecal concentration of 20 µM^42,43^, 24 h before DSS treatment, and was kept in the luminal perfusion circuit during the subsequent DSS exposure. Light microscopic analysis revealed that LCA preserved villus-like structures in the presence of DSS (**Figure 3A**) and prevented the DSS-induced decline of villus height (**Figure 3B**). Further, LCA treatment stabilized epithelial barrier integrity, reflected by reduced DSS-induced permeability, comparable to tissue models without DSS treatment (**Figure 3C**).

**Figure 3:**
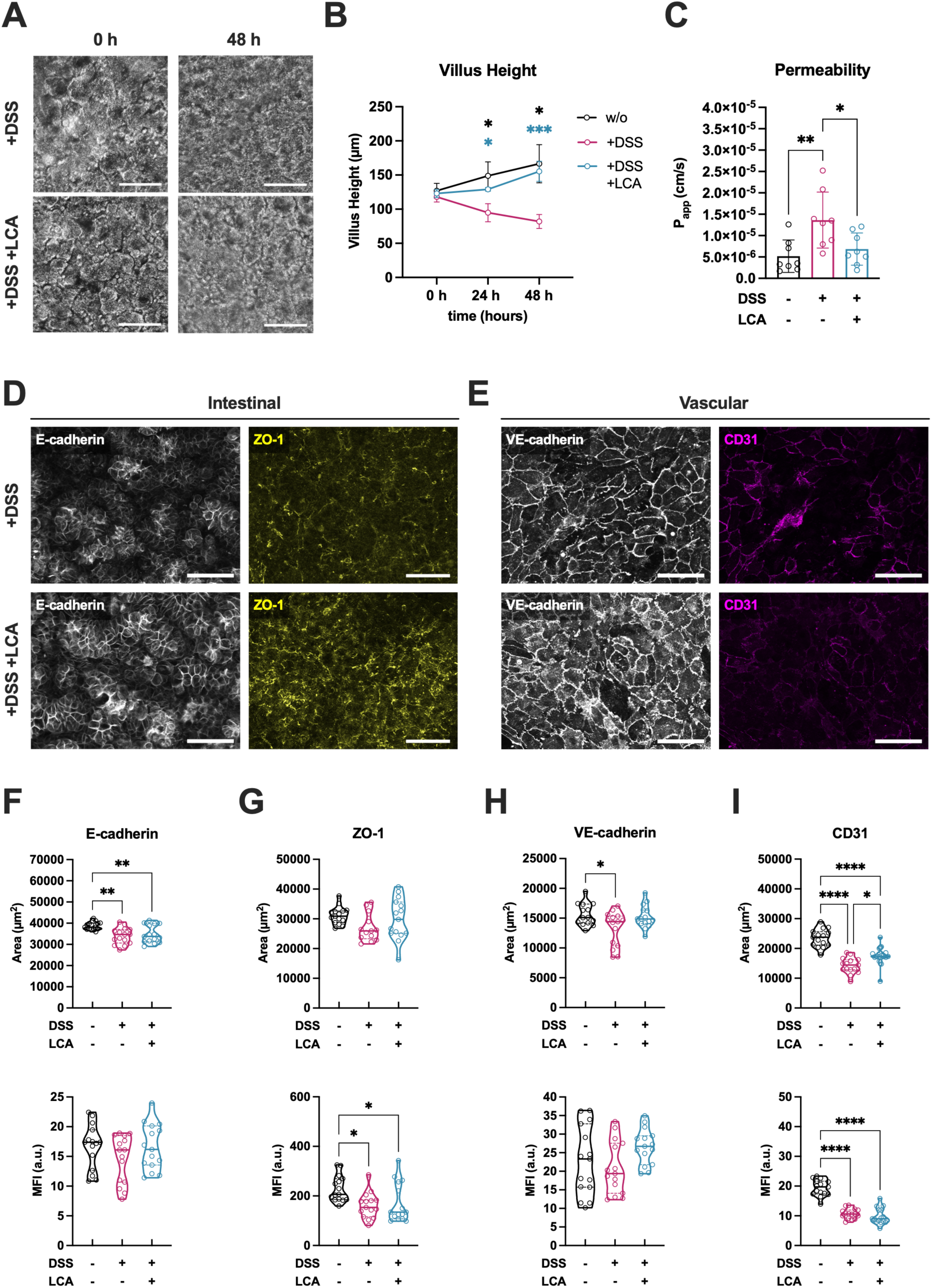
LCA prevents DSS-induced morphological alterations and barrier disruption. **(A)** Light microscopic monitoring of untreated models (w/o) and models treated with DSS (+DSS) or DSS/LCA (+DSS +LCA) for 48 h. **(B)** Measurement of villus height in untreated models (w/o) and models treated with DSS (+DSS) or DSS/LCA (+DSS +LCA) from 0 h to 48 h of treatment. Data points represent mean ± SD of 4 independent experiments (n = 4) with 3 individual MDM donors. *p ≤ 0.05, ***p ≤ 0.001 (Two-way ANOVA with Tukey’s multiple comparisons test). **(C)** Permeability of untreated models (w/o) and models treated with DSS (+DSS) or DSS/LCA (+DSS +LCA) after 48 h. Bars represent mean ± SD of 8 independent experiments (n = 8) with 6 individual MDM donors. *p ≤ 0.05, **p ≤ 0.01 (Paired two-tailed t test). **(D,E)** Representative fluorescence images of intestinal **(D)** and vascular cell layers **(E)** from models with DSS (+DSS) or with DSS/LCA (+DSS +LCA) treatment after 48 h. Caco-2 cells were stained for E-cadherin (white) and ZO-1 (yellow). HUVECs were stained for VE-cadherin (white) and CD31 (magenta). Scale bars, 100 µm. **(F-I)** Quantification of covered area (upper row, in µm^2^) and mean fluorescence intensities (lower row, MFI) of E-cadherin **(F)**, ZO-1 **(G)**, VE-cadherin **(H)**, and CD31 **(J)** signals. Violin plots with indicated median and quartiles of 3 independent experiments (n = 3) with 3 individual MDM donors. 5 different images were taken for each experiment. *p ≤ 0.05, **p ≤ 0.01, ****p ≤ 0.0001 (One-way ANOVA with Tukey’s multiple comparisons test).

Quantitative analysis of immunofluorescence images further supported the protective effect of LCA. Treatment with LCA partially restored the MFI and covered area of AJCs on both the intestinal epithelium and the vascular endothelium (**Figure 3D,E**). In the intestinal compartment, LCA induced a trend of increase in E-cadherin and ZO-1 expression indicating a positive effect on maintaining epithelial junctional integrity (**Figure 3F,G**). Further, LCA induced a modest increase in VE-cadherin MFI and coverage, and a significant enhancement of CD31 expression, implicating improved endothelial junctional continuity under DSS-induced inflammatory conditions (**Figure 3H,I**).

### 2.4 Vascular and intestinal inflammation is reduced by LCA

Due to the beneficial effects of LCA on intestinal morphology, barrier function, and AJCs (**Figure 3**), its immunomodulatory role within intestinal and vascular cell compartments was further investigated (**Figure 4**). Notably, a significant reduction of IL-6 (**Figure 4A**) and IL-8 (**Figure 4C**) was observed following LCA administration at 24 h in the vascular supernatant. Additionally, TNF-α concentrations tended to be reduced in response to LCA treatment after both 24 h and 48 h (**Figure 4B**).

**Figure 4:**
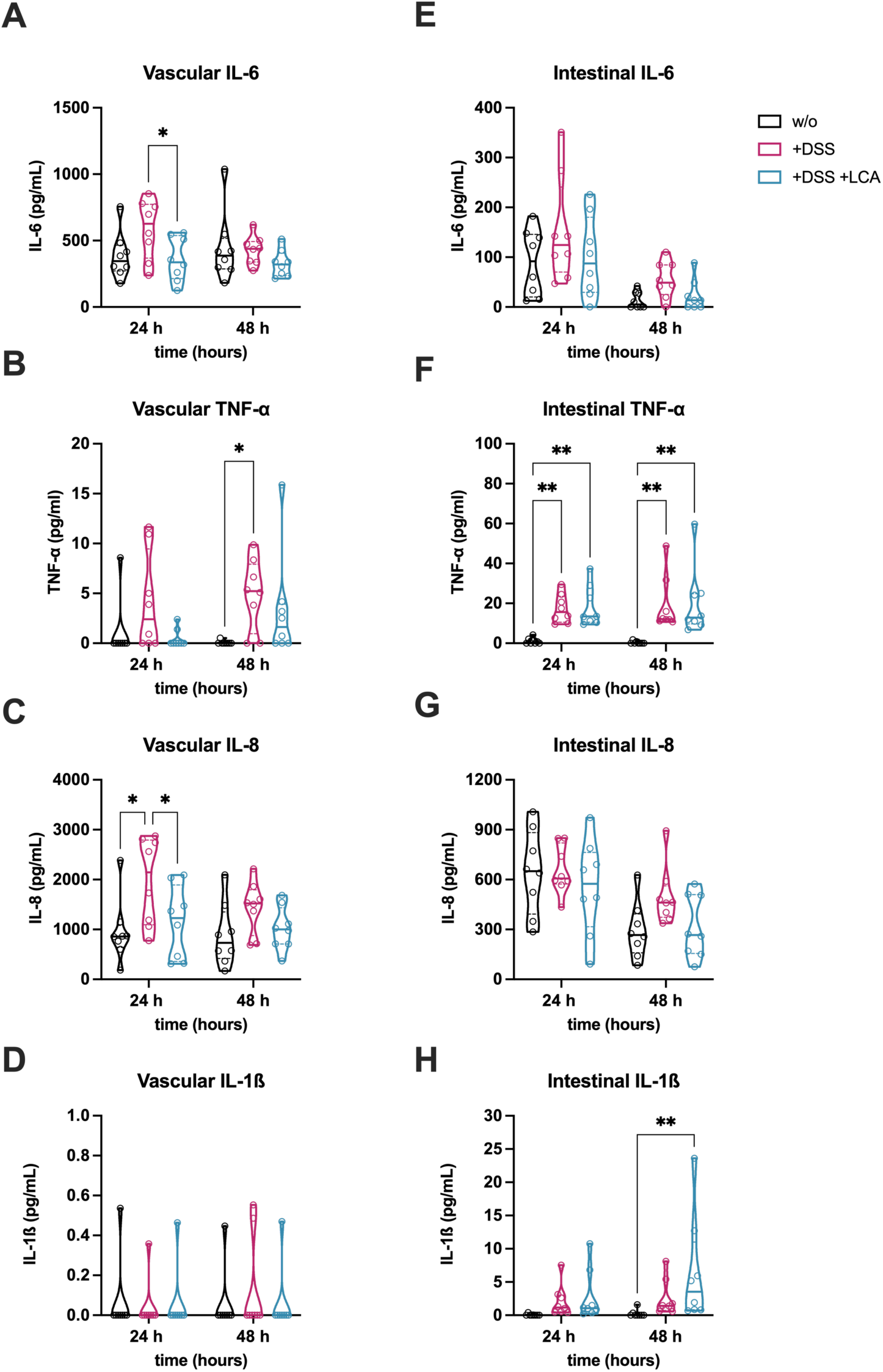
LCA reduces inflammation in the colitis-on-chip model. Release of pro-inflammatory cytokines (IL-6, TNF-α, IL-8, IL-1β) in vascular **(A-D)** and intestinal **(E-H)** medium supernatants of untreated models (w/o) and models treated with DSS (+DSS) or DSS/LCA (+DSS +LCA). Violin plots with indicated median and quartiles of 8 independent experiments (n = 8) with 8 individual MDM donors. *p ≤ 0.05, ***p ≤ 0.001, ****p ≤ 0.0001 (Two-way ANOVA with Tukey’s multiple comparisons test).

No significant differences were detected for IL-1ß (**Figure 4D**), likely due to cytokine concentrations near the detection limit. In intestinal supernatants, LCA treatment showed a trend of reduced IL-6 and IL-8 levels after 48 h (**Figure 4E,G**). In contrast, LCA treatment did not reduce TNF-α levels (**Figure 4F**) and was associated with an increase in IL-1ß compared to untreated models after 48 h (**Figure 4H**). Overall, LCA modulated vascular cytokine levels to a greater extent than those in the intestinal compartment, potentially due to enhanced barrier function and reduced translocation of luminal LPS and higher abundance of MDMs residing in the vascular cell layer.

To characterize the contribution of MDMs to inflammatory responses, cytokine levels were compared from models with and without MDMs. The release of IL-6, TNF-α, and IL-8 was dependent on the presence of integrated MDMs in the vascular channel (**Figure 5A-C**), associated with increased release of IL-6 into the luminal side of the model (**Figure 5D**). In contrast, no differences in the cytokine profile with or without MDMs were found for intestinal TNF-α (**Figure 5E**).

**Figure 5:**
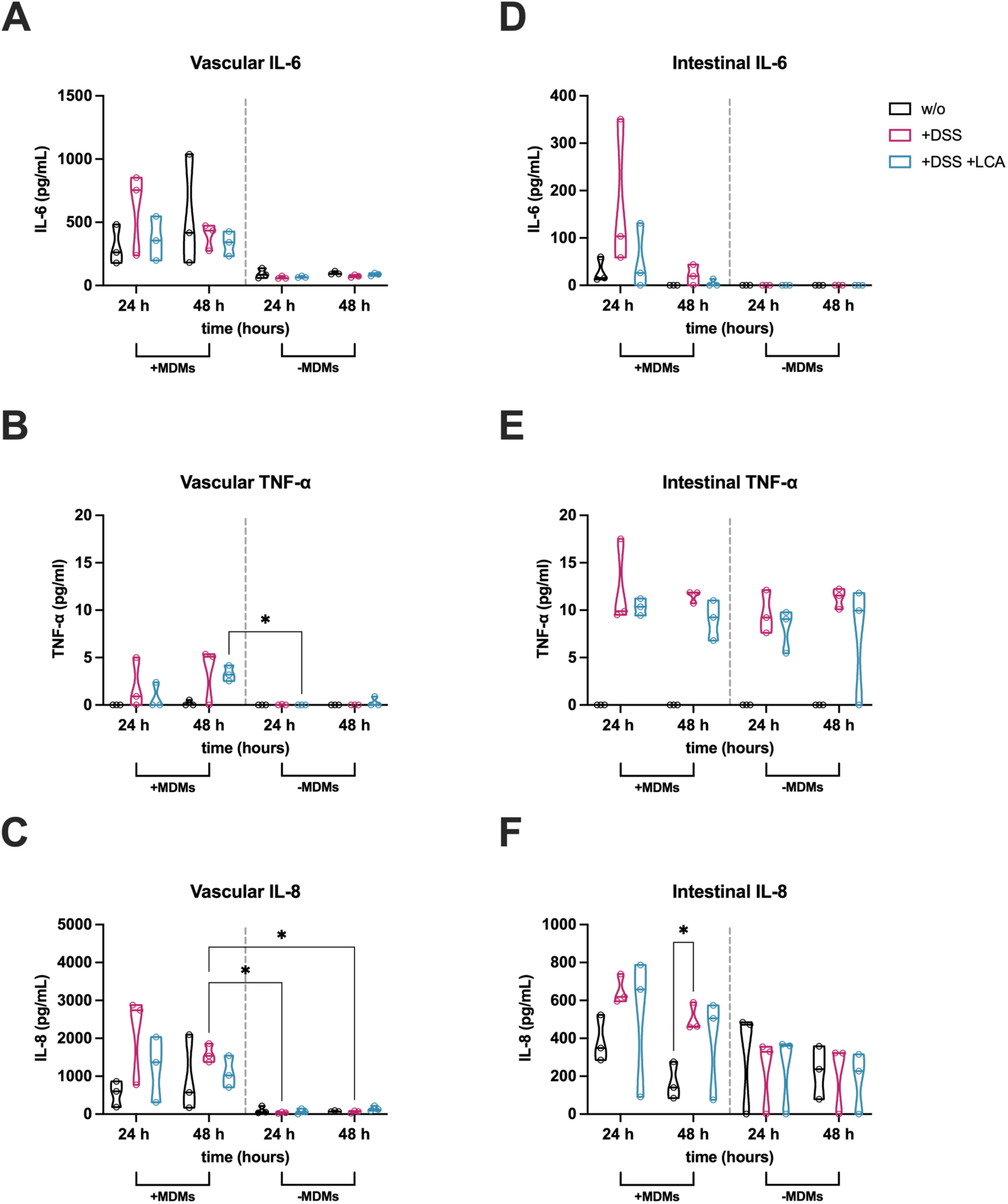
Pro-inflammatory cytokine release is partially dependent on the presence of MDMs. Measurement of cytokine levels from vascular **(A-C)** and intestinal **(D-F)** medium supernatants of untreated models (w/o) and models treated with DSS (+DSS) or DSS/LCA (+DSS +LCA) either with (+MDMs) or without (-MDMs) MDMs (separated by grey dotted line). Violin plots with indicated median and quartiles of 3 independent experiments (n = 3) with 3 individual MDM donors. *p ≤ 0.05 (Two-way ANOVA with Tukey’s multiple comparisons test).

### 2.5 Application of LCA prevents DSS-induced restriction of cell proliferation

Previous studies in mice reported a restricted intestinal epithelial proliferation induced by DSS treatment^44,45^. It was investigated whether LCA can support proliferation in the presence of DSS based on an expression of the proliferation marker Antigen Kiel 67 (Ki-67). The analysis of immunofluorescence images showed a decrease in Ki-67 signals upon DSS treatment (**Figure 6A**), which was to some extent rescued by LCA. Automated image quantification (**Supplementary Figure S6**) confirmed that DSS treatment induced a significant reduction in Ki-67 MFI in intestinal epithelial cells (**Figure 6B**). LCA treatment significantly improved epithelial cell proliferation compared to models treated with DSS only, implicating a preservation of intestinal cell proliferation. However, LCA treatment did not fully maintain proliferation levels that were observed in untreated models.

**Figure 6:**
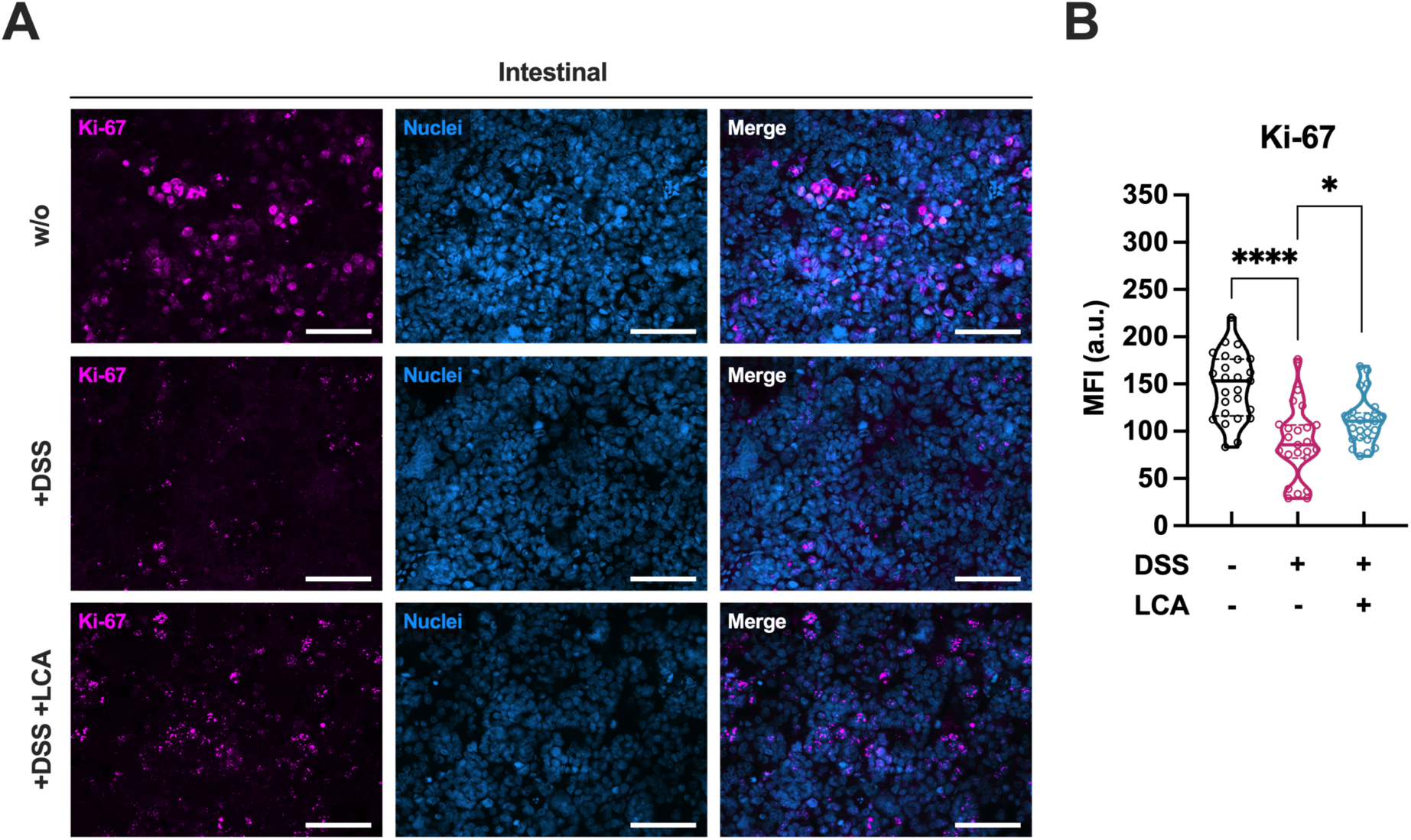
LCA prevents the disruption of intestinal cell proliferation in the colitis-on-chip model. **(A)** Representative immunofluorescence images of intestinal cell layers from untreated models (w/o) and models treated with DSS (+DSS) or DSS/LCA (+DSS +LCA) after 48 h. Caco-2 were stained for Ki-67 (magenta) and DAPI (nuclei, blue). Scale bars, 100 µm. **(B)** Quantification of Ki-67 mean fluorescence intensity (MFI) within the DAPI-positive nuclei. Violin plots with indicated median and quartiles of 3 independent experiments (n = 3) with 3 individual MDM donors. 5 different images were taken for each experiment. *p ≤ 0.05, ****p ≤ 0.0001 (Unpaired two-tailed t test).

### 2.6 LCA differentially acts on FXR and TGR5 to modulate barrier function and immune responses

Next, the role of the nuclear FXR and the transmembrane TGR5 in the observed amelioration of the DSS-induced tissue damage was investigated. These receptors are known to bind LCA with high affinity^46–48^ and were suggested to play a potentially protective role in mice^49,50^. However, receptor antagonists guggulsterone targeting FXR and SBI-115 acting on TGR5 showed a dose-dependent toxicity on HUVECs (**Supplementary Figure S7A**) and Caco-2 cells (**Supplementary Figure S7D**). Therefore, concentrations of 3 µM for guggulsterone and 1 µM for SBI-115 were selected, which did not interfere with cell viability up to 72 h of treatment and have been reported to inhibit FXR^51,52^ or TGR5^53^ activity effectively. Blocking FXR activity with guggulsterone interfered with the protective effect of LCA treatment on DSS-induced barrier permeability (**Figure 7A**), whereas no significant impact was observed for SBI-115 targeting TGR5. Quantification of Ki-67 expression revealed that in contrast to SBI-115, which had no effects, guggulsterone treatment mediated a substantial depression of Ki-67 in the colitis-on-chip model treated with LCA (**Figure 7B**). However, none of the antagonists had an effect on IL-6 and vascular IL-8 levels (**Supplementary Figure S8).** Nevertheless, a significant increase in intestinal IL-8 levels was observed after treating with 1 µM SBI-115 (**Supplementary Figure S8D**), demonstrating a potential role of the TGR5 receptor in partially modulating LCA-mediated intestinal IL-8 responses.

**Figure 7:**
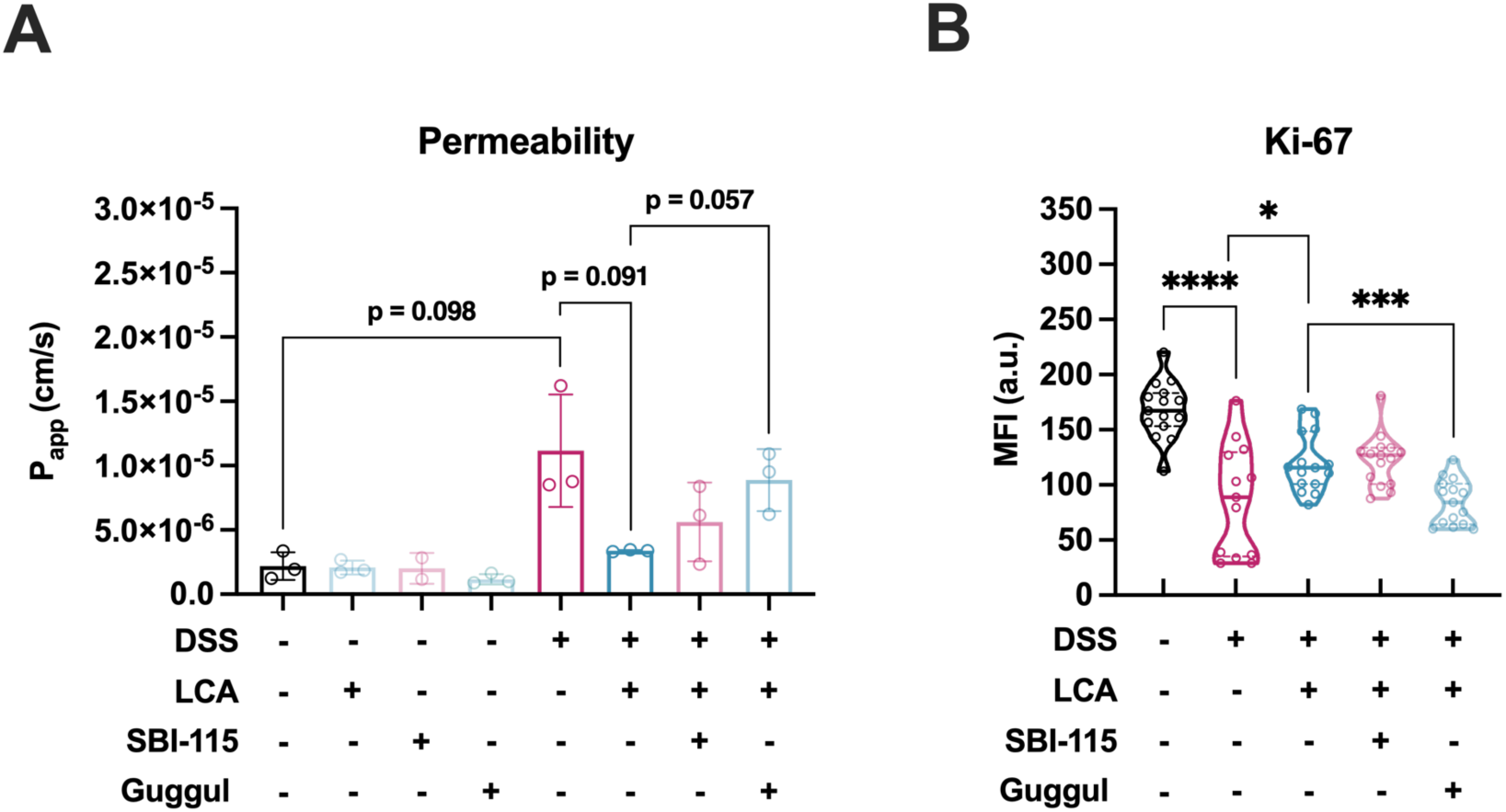
LCA acts via FXR to maintain barrier integrity and cell proliferation. **(A)** Measurement of FITC-dextran permeability (shown as calculated permeability coefficient P_app_ in cm/s) after 48 h. Bars represent mean ± SD of 3 independent experiments (n = 3) with 3 individual MDM donors. Significances are directly indicated as written p-values (Paired two-tailed t test). One data point (GMM +LPS +1 µM SBI-115) was excluded due to leakage of the biochip. **(B)** Quantification of Ki-67 mean fluorescence intensity (MFI) within the identified nuclei in untreated models (w/o) or models treated with DSS (+DSS), DSS/LCA (+DSS +LCA), DSS/LCA/SBI-115 (1 µM) (+DSS +LCA +SBI-115), and DSS/LCA/guggulsterone (3 µM) (+DSS +LCA +Guggul). Violin plots with indicated median and quartiles of 3 independent experiments (n = 3) with 3 individual MDM donors. *p ≤ 0.05, ***p ≤ 0.001, ****p ≤ 0.0001 (Unpaired two-tailed t test).

## 3. Discussion

IBD is a multifaceted disease involving complex interactions across multiple cellular and molecular pathways and mechanisms. Its complexity highlights the importance of more reliable experimental *in vitro* models recapitulating relevant clinical disease features under near-physiological conditions.

This study showcases a microfluidic DSS-induced colitis-on-chip model that effectively recapitulates most of the colitis pathophysiological features that are observed in humans^54^ and mice^25^. Thus, the model effectively reproduced important disease hallmarks, such as intestinal epithelial barrier permeability, disruption of AJC networks, pro-inflammatory cytokine release, villus atrophies, and reduction of intestinal cell proliferation, as it was recently shown in comparable DSS-induced IBD chip models^37,38^. MDMs integrated in the model were shown to be the main driver of vascular inflammation, whereas contributing to a lesser extent to the cytokine responses in the intestinal compartment. The administration of DSS in combination with LPS further appeared to mediate direct cytokine responses in Caco-2 or HUVECs, as the cytokine levels were not significantly altered for TNF-α and IL-1ß in the absence of MDMs. This might implicate a role of both Caco-2 and HUVECs in releasing TNF-α and IL-1ß and contributing to inflammatory responses upon stimulation^55,56^. These findings highlight the potential of OoC technology in dissecting cell-specific responses by selectively tuning the composition of the model.

Leveraging this colitis-on-chip model, the impact of LCA in protecting against the DSS-induced inflammation and the associated barrier disruption was explored. Since it is well described that LCA is a bacterial-derived metabolite that exerts immunomodulating activity and can promote intestinal barrier integrity, it was investigated whether there is a causal link between the reduction of LCA observed in clinical patients and the onset of IBD. The colitis-on-chip model was used to explore the effects of LCA on protecting against inflammation-induced DSS, which disrupts the intestinal barrier. Given LCAs well-established function for its immunomodulating properties^20,21,27,28,57^, its role in maintaining intestinal barrier integrity upon DSS stimulation was investigated. At human-relevant physiological concentrations, LCA attenuated the colitis phenotype by preserving the intestinal barrier function, AJC integrity, and 3D villus morphology. This was likely due to LCA-mediated improvement of self-renewal by stimulating intestinal epithelial cell proliferation under inflammatory conditions caused by DSS treatment.

By applying SBI-115 as a TGR5 antagonist and guggulsterone as a FXR antagonist, it was investigated whether LCA-mediated protective effects could be reversed by inhibiting these BA receptors. It was demonstrated that LCA acts via FXR to mediate protective functions on barrier integrity and cell proliferation. Upon binding of LCA, FXR translocates into the nucleus and regulates the transcription of target genes^58^. The preservation of barrier function might be mediated via the FXR-dependent inhibition of the myosin light chain kinase (MLCK) pathway, leading to the improvement of AJCs such as ZO-1, occluding, and claudin-1, as it was previously described for the SBA chenodeoxycholic acid^59^. In accordance with our observations in this study, inhibiting the TGR5 pathway had no effects on the LCA-mediated barrier preservation. Furthermore, taurodeoxycholic acid has been shown to alleviate LPS-induced intestinal injury in mice by acting on the FXR receptor and increasing cellular myelocytomatosis oncogene (c-Myc)-dependent enterocyte proliferation^60^. These results align with the findings for LCA in the colitis-on-chip model, which point out a role of FXR in ameliorating intestinal permeability.

In mice, it was shown that the semi-synthetic agonist obeticholic acid (INT-747) was protective against loss of epithelial integrity and associated ulceration, immune cell infiltration, and pro-inflammatory cytokine release in chemically-induced colitis models using stimulation with DSS or 2,4,6-trinitrobenzene sulfonic acid (TNBS)^61^. Also the FXR agonist 6α-ethyl-3α,7α,23-trihydroxy-24-nor-5β-cholan-23-sulfate (INT-767), which targets TGR5 at the same time was protective for a colitis phenotype in a DSS mouse model^62^. Further, the TGR5 agonist OM8 was found to alleviate colitis in mice by maintaining tight junction protein expression and preventing intestinal epithelial cell apoptosis^63^.

The findings of this work emphasize the role of LCA as an important mediator of the intestinal homeostasis. However, regarding its potential therapeutic use, the time point of its administration in IBD is critical. Previous studies have indicated contradictory effects of probiotic or postbiotic treatment upon active IBD characterized by a highly permeable intestinal epithelial barrier and excessive inflammation. In this context, the administration of probiotic bacteria in a DSS-induced colitis model^37^ or postbiotic SCFAs in a gut-liver model^64^ where an active IBD phenotype was present resulted in excessive inflammation, presumably through translocation into the adjacent vascular perfusion and following immune cell stimulation. These findings suggest that the timing of application in severe IBD flares could cause undesirable adverse events such as inflammatory immune cell activation or vascular injury. It indicates that postbiotics such as LCA may rather have a preventive effect before the onset or during the early stages of IBD and for maintaining disease remission. Future studies should therefore examine the effects of postbiotic administration during different IBD stages.

Although the colitis-on-chip model shows promising potential as a model of IBD, some limitations need careful consideration for its future applications. While Caco-2 cells are widely used for modeling intestinal permeability and transporter function, they lack key metabolic enzymes such as cytochrome P450 3A4 (CYP3A4), limiting their utility for studying drug metabolism and complex host responses compared to more physiologically relevant organoid-derived enterocytes^65^. Integrating intestinal organoid-derived cells collected from IBD patients and healthy donors and their use in an immunocompetent chip-based intestinal model may reduce the need for exogenous compounds such as DSS. Recently, it was demonstrated that epithelial tissue samples isolated from both CD and UC patients can be expanded as intestinal organoids and subsequently cultured in a microfluidic chip platform for up to 11 days, resulting in the recapitulation of disease-specific phenotypes and responses^66^. This approach could potentially provide physiologically relevant models for studying microbiota-based therapeutic methods in a personalized manner.

Despite recapitulating important hallmarks of experimental colitis, including the disruption of AJCs, associated loss of barrier function, and reduced cell proliferation^67,68^, the colitis model rather reflects an acute and unselective IBD-like phenotype without addressing genetic and environmental components and the chronic refractory nature of the disease^69^.

Combining a model based on patient-derived cells with multi-omics analyses from clinical samples such as serum, feces, and tissue biopsies will pave the way for connecting alterations in the microbial metabolome pool observed in IBD patients and disease pathophysiology to more specifically screen individual metabolites of interest^2^.

Although the model incorporates a significant level of complexity with the inclusion of MDMs as an integral component of immune cells, it currently falls short of fully capturing the plasticity and diversity of human immune cell populations. This is particularly true for T cells, which include T helper (Th) cells (CD4^+^), cytotoxic T cells (CD8^+^), and regulatory T cells (Treg), along with their contributions to the IBD phenotype. Differences in CD and UC have been shown to arise from distinct Th cell responses and unique cytokine patterns^70^. Thus, including key disease drivers such as Th1/Th17 cells for CD^71^ and Th2 cells for UC^72^ into the IoC model could provoke cytokine responses that better reflect the *in vivo* situation.

Since LCA can act on various BA receptors^73^, there might be other receptor targets than anticipated in this study. For example, LCA has been shown to modulate Th1 activation through the vitamin D receptor^74^. Moreover, metabolites derived from LCA, such as 3-oxolithocholic acid (3-oxoLCA) and isolithocholic acid (isoLCA), have recently been identified to influence T cell differentiation by inhibiting the retinoic receptor-related orphan nuclear receptor γt (RORγt)^75^. Specifically, they suppress the differentiation of CD4^+^ T cells into Th17 cells. Since inflammatory Th cell populations are implicated as drivers of intestinal inflammation in IBD, targeting their differentiation and polarization could be a promising strategy to restore the immune balance of the host.

Furthermore, integrating a living microbiota or relevant microbial communities will be particularly important in future studies. Several studies have already highlighted direct antimicrobial actions of LCA on human pathogens and pathobionts, such as *C. difficile*^76,77^ or *C. albicans*^78^, microbes that are commonly enriched in IBD patients^79,80^. The application of LCA could modulate the host resilience and directly influence the growth, adhesion, and invasion of these intestinal microbes. Integrating living bacteria could shed more light on host-microbiota crosstalk that is altered during IBD and how selective application of postbiotics or other microbial-derived products shape these interactions.

In summary, the colitis-on-chip model emulates central hallmarks of the human colitis phenotype and allows for screening of microbiota-derived metabolites, such as LCA, to investigate their therapeutic effect for clinical translation. As a model with scalable complexity and simple controllability, it can be particularly useful to emulate relevant disease characteristics in a human-based manner and to dissect underlying disease-driving mechanisms and causal relationships between microbiota and the host.

## 4. Materials and methods

### 4.1 Ethics statement

Human peripheral blood was collected from consenting and informed healthy volunteers in accordance with the Declaration of Helsinki guidelines. The collection and usage of blood samples for this study were approved by the institutional ethics committee of the Jena University Hospital (permission number 2207-01/08). HUVECs were isolated under ethical approval 2020-1684, 3939-12/13 following written informed consent from donors.

### 4.2 Cell culture

All cell culture incubation procedures were performed at 37°C and 5% CO_2_ in a humidified cell culture incubator, unless stated otherwise. Cells used in this study have been tested for mycoplasma contaminations using a PCR mycoplasma test kit (Minerva Biolabs, Berlin, Germany) following the instructions of the manufacturer.

Caco-2 epithelial cells (accelerate GmbH, Hamburg, Germany) were thawed and seeded in culture flasks at a density of 0.8 × 10^4^ cells/cm^2^. Cells were cultured in DMEM with 4.5g/L glucose (Lonza, Cologne, Germany) supplemented with 10% fetal bovine serum (FBS, Capricorn Scientific, Ebsdorfergrund, Germany), 1 mM sodium pyruvate solution (Capricorn), 1× MEM non-essential amino acids (Capricorn), 5 mg/mL holo-transferrin (Merck, Darmstadt, Germany) and 20 µg/mL gentamycin (Merck). Medium was exchanged every three days until further passaging or seeding into biochips.

HUVECs were isolated, expanded, and cryopreserved as previously described^81^. Cryopreserved HUVECs were thawed and seeded in culture flasks at a density of 1.3 × 10^4^ cells/cm^2^. HUVECs were cultured in Endothelial Cell Growth Medium (ECGM) MV (Promocell, Heidelberg, Germany) including supplements and 1× antibiotic antimycotic solution (AAS, Merck). Medium was exchanged every two days until further passaging or seeding into biochips. HUVECs were used up to passage 5.

Human peripheral blood was drawn from healthy donors and either collected in EDTA tubes (Sarstedt, Nümbrecht, Germany) for PBMC isolation or serum gel tubes (Sarstedt) to obtain autologous donor serum for medium supplementation. All donors were informed about the study and gave written consent. All procedures were performed according to the approved guidelines and regulations and to the guidelines set forth in the Declaration of Helsinki. PBMC isolation and serum extraction were performed as previously described^34,82^. Briefly, PBMCs were isolated from donor blood by density gradient centrifugation using lymphocyte separation medium (Capricorn). PBMCs were seeded at a density of 1 × 10^6^ cells/cm^2^ in X-VIVO 15 medium (Lonza) with 10% autologous serum, 10 ng/mL macrophage colony-stimulating factor (M-CSF), 10 ng/mL granulocyte-macrophage colony-stimulating factor (GM-CSF) and 1× AAS. Cells were cultured for five days with medium exchange before seeding into biochips.

### 4.3 Biochip fabrication

Biochips type BC002 were manufactured by Dynamic42 GmbH (Jena, Germany) from injection-molded polybutylene terephthalate bodies. The biochip is composed of two independent cavities for cell culture. Each biochip cavity consists of a top (area: 2.18 cm^2^) and a bottom (area: 1.62 cm^2^) culture channel, separated by a polyethylene terephthalate (PET) membrane (TRAKETCH Sabeu, Radeberg, Germany). The PET membrane is 12 µm thin and consists of 1 × 10^5^ pores/cm^2^ with a pore diameter of 8 µm. The top channel has a total volume of 290 µL and the bottom channel of 270 µL, including adjacent channels, inlets, and outlets. Cells are seeded in the top channel in 200 µL volume, while 150 µL cell suspension is introduced in the bottom channel. Prior to the perfusion, microfluidic medium reservoirs were attached to the biochips. Biochips were connected to a peristaltic pump system (Masterflex, VWR International, Darmstadt, Germany) with platinum-cured 2-stop silicone tubing (Dynamic42 GmbH).

### 4.4 Intestinal model assembly

Biochips were first sterilized with 70% ethanol (VWR) and then washed with AQUA AD iniectabilia (B.Braun, Melsungen, Germany) and phosphate buffered saline (PBS, Lonza). Biochip membranes were coated in the top channel with 50 µg/mL bovine collagen A (PAN-Biotech, Aidenbach, Germany). HUVECs were seeded in the top channel, at a density of 1.38 × 10^5^ cells/cm^2^ in supplemented ECGM MV with 1× AAS. HUVECs were cultured for 48 h with daily medium exchange under static conditions in the biochips. Prior to the seeding of MDMs, the medium in the top channel was changed to vascular perfusion medium (VPM) consisting of supplemented ECGM MV, 10% autologous donor serum, 10 ng/mL M-CSF and GM-CSF and 1× AAS. MDMs were seeded at a density of 0,45 × 10^5^ cells/cm^2^ on top of the HUVECs. After 24 h, membranes were coated on the other side via the lower channel with 50 µg/mL bovine collagen A. Caco-2 cells were seeded on collagen-coated membranes in the bottom channel at a density of 2.16 × 10^5^ cells/cm^2^ in intestinal seeding medium containing DMEM with 4.5 g/L glucose supplemented with 20% FBS, 1× MEM non-essential amino acids (Capricorn), 1 mM sodium pyruvate solution (Capricorn), 5 mg/mL holo-transferrin (Merck) and 20 µg/mL gentamycin (Merck). Biochips were incubated upside down for 24 h to facilitate attachment of the Caco-2 cells to the membrane. The intestinal medium was changed to intestinal perfusion medium consisting of intestinal seeding medium with only 10% of FBS. Medium exchange was also performed in the vascular channel, with vascular perfusion medium. Following complete static assembly, the models were dynamically perfused in both channels at 50 µL/min equaling shear stress rates of 0.013 dyn/cm^2^ (0.0013 Pa) in the top channel and 0.006 dyn/cm^2^ (0.0006 Pa) in the bottom channel^34^. Intestinal models were pre-perfused for 72 h prior to the IBD-like disease induction.

### 4.5 IBD-like phenotype induction and treatments

Medium was exchanged in the vascular and top channel after pre-perfusion for 72 h. The medium of the intestinal channel was replaced and 100 ng/mL lipopolysaccharide (LPS, CAS number: 93572-42-0, from *E. coli* O111:B4, Merck) was added to the intestinal tissue. The colitis-like phenotype was induced after 24 h of pre-stimulation with LPS by adding 1.5% DSS (36-50 kDa, CAS number: 9011-18-1, MP Biomedicals, Eschwege, Germany) and 100 ng/mL LPS into the intestinal perfusion medium for 48 h with redosing after 24 h. Untreated models were perfused with intestinal perfusion medium containing 100 ng/mL LPS without DSS.

Lithocholic acid (LCA, CAS number: 434-13-9, Merck) was solubilized in dimethyl sulfoxide (DMSO, Merck) to a stock concentration of 20 mM. The stock was diluted 1:1000 in culture medium to a final concentration of 20 µM (0.1% DMSO). LCA was administered in combination with LPS 24 h prior to the application of DSS and was administered every 24 h until analysis.

TGR5 antagonist SBI-115 (CAS number: 882366-16-7, MedChemExpress, Sollentuna, Sweden) was solubilized in DMSO to a stock concentration of 100 mM. FXR antagonist guggulsterone (CAS number: 95975-55-6, MedChemExpress) was solubilized in DMSO to a stock concentration of 50 mM. Both antagonists were further diluted in culture medium to the respective test concentration without exceeding DMSO concentrations of 0.1%. SBI-115 or guggulsterone were applied in combination with LPS and LCA 24 h prior to the application of DSS and were administered every 24 h until analysis.

2D plate experiments were performed for excluding LCA, SBI-115, and guggulsterone toxicity on both Caco-2 and HUVECs. Therefore, Caco-2 were plated at a density of 6 × 10^3^ cells/cm^2^ and cultured for two weeks in intestinal perfusion medium with medium exchange every three to four days. HUVECs were plated at a density of 1.6 × 10^4^ cells/cm^2^ and were cultured for 24 h. Subsequently, cells were treated for up to 72 h with LCA, SBI-115, or guggulsterone with daily redosing.

### 4.6 Cell viability assay

Cell viability was determined by using the CellTiter-Glo Luminescent Cell Viability Assay (Promega, Walldorf, Germany). Viability was measured in 2D plate experiments and 3D IoC models. Briefly, wells or chip channels were washed with PBS with calcium and magnesium (w Ca/Mg, +/+). For plate experiments, 50 µL of PBS +/+ was added to each well. Biochip membranes were excised by using a scalpel and transferred to a 48-well microplate containing 100 µL PBS +/+. CellTiter-Glo Reagent was added in a 1:1 ratio to either the plate wells or the excised biochip membranes. Cell lysis was performed for 2 min on an orbital plate shaker and the plate was subsequently incubated for 10 min at room temperature in the dark. The solution was transferred to a white 96-well plate (LUMITRAC 600, Greiner, Frickenhausen, Germany) and the luminescence was measured in a microplate reader (Infinite 200 PRO, Tecan, Crailsheim, Germany).

### 4.7 Permeability assay

Permeability of IoC models was determined after diffusion of fluorescein isothiocyanate (FITC-dextran, 3-5 kDa, CAS number: 60842-46-8, Merck) from the intestinal channel to the vascular channel. Cell culture medium in both channels was replaced by prewarmed phenol red-free William’s Medium E (PAN-Biotech). FITC-dextran was diluted in phenol red-free William’s Medium E to a stock concentration of 2 mg/mL. A volume of 250 µL of the FITC-dextran stock solution was added two times in the lower intestinal channel to minimize dilution effects. All inlets and outlets of the biochips were closed and chips were incubated upside down in the dark at 37°C and 5% CO_2_ for 1 h. Supernatants from both intestinal and vascular channels were collected and transferred to a black 96-well microplate (Greiner). Fluorescence intensities were measured in a microplate reader (Infinite 200 PRO, Tecan) at 488 nm excitation and 520 nm emission. FITC-dextran concentrations were determined from the standard curve. The apparent permeability coefficient was calculated from permeated FITC-dextran concentrations using the following equation (1)^37^:

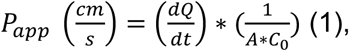

where *dQ/dt* is the steady-state flux within the incubation time (μg/s), A is the surface area of cultivated cells on the membrane (cm^2^), and C_0_ is the initial concentration of the FITC-dextran stock solution (μg/cm^3^).

### 4.8 Measurement of villus height

The morphology of the IoC models was observed and monitored using a phase contrast inverted cell culture microscope (ZEISS Primovert, Carl Zeiss AG, Jena, Germany). Villus height was determined by measuring the Z-position between the surface of the porous membrane facing the lowest position of the intestinal epithelial cells and the villus tip (five randomly selected areas per condition and time point).

### 4.9 Cytokine measurement

Medium supernatants were collected from vascular and intestinal reservoirs every 24 h. Supernatants were collected, centrifuged to remove remaining cell debris, and stored at −80°C for further analysis. Cytokine concentrations were measured by using the ELISA MAX Deluxe Set Human for IL-6, IL-1β, TNF-α, and IL-8 (BioLegend, Amsterdam, The Netherlands). The assay was performed according to the provided protocol from the manufacturer. The volumes of samples and assay reagents specified by the manufacturer were reduced by half. For the measurement of IL-6 and IL-8, vascular and intestinal samples were pre-diluted at a 1:2 ratio in assay buffer. IL-1β and TNF-α levels were determined from undiluted supernatants. Absorption was measured in a microplate reader (Infinite 200 PRO, Tecan) at 450 nm with a reference wavelength of 570 nm. Cytokine concentrations were calculated from the plotted standard curve of each assay.

### 4.10 Immunofluorescence staining

Biochips were disconnected from the perfusion system. Both the vascular and intestinal channels were washed with cold PBS +/+. Cells were fixed within the biochips with ice-cold methanol (Carl Roth, Karlsruhe, Germany) at −20°C for 15 min. Cells were washed three times with PBS +/+. Membranes with fixed cells were recovered from biochips by excision with a scalpel. Membranes were incubated in permeabilization/blocking solution containing PBS +/+ with 3% normal donkey serum (Abcam, Amsterdam, The Netherlands) and 0.1% saponin (Carl Roth). Epithelial cells were stained with primary antibodies mouse anti-E-cadherin (2.5 µg/mL, BD Bioscience, Heidelberg, Germany), rabbit anti-ZO-1 (1.25 µg/mL Thermo Fisher Scientific), or mouse anti-Ki-67 (2.5 µg/mL, BD Bioscience) and vascular cells for mouse anti-CD31 (0.5 µg/mL, Cell Signaling Technology, Leiden, The Netherlands), rabbit anti-CD68 (2.92 µg/mL Cell Signaling Technology), and goat anti-VE-cadherin (2 µg/mL, Biotechne, Nordenstadt, Germany) in permeabilization/blocking solution at 4°C overnight. After incubation, membranes were washed with PBS +/+/ 0.1% saponin and incubated with secondary antibodies DAPI, donkey-anti-mouse AF555, donkey-anti-mouse AF647, donkey-anti-rabbit-AF488, donkey-anti-rabbit-Cy3 (Jackson ImmunoResearch, St. Thomas’ Place, United Kingdom, 7.5 µg/mL), or donkey-anti-goat-AF647 (all Thermo Fisher Scientific, 10 µg/mL) at room temperature for 1 h. Membranes were washed afterwards and mounted in fluorescence mounting medium (Agilent, Waldbronn, Germany) on microscopic glass slides.

### 4.11 Image acquisition and analysis

Fluorescence images were acquired using the ZEISS Axio Observer Z1 with ApoTome.2 or ZEISS Axio Observer 7 with Apotome 3 (Carl Zeiss AG) and Plan Apochromat 20x/0.8 M27 objective. Images were taken as Z-stacks. Z-stack images were Apotome Raw converted and merged as orthogonal projection. 3D reconstructions were generated from Z-stack images using the 3D module in the ZEISS ZEN 2 Pro software (Carl Zeiss AG). The background noise was removed by applying a Gaussian filter.

Five images of randomly selected areas of each condition were taken for subsequent image analysis. Quantification of fluorescence images were performed using the analysis software CellProfiler (Broad Institute, Cambridge, MA, USA)^83^.

Junctional networks were extracted from intestinal E-cadherin and ZO-1 and vascular VE-cadherin and CD31 fluorescence signals. The images were processed in the “Morph” module with the “openlines” operation to enhance linear structural elements and to suppress scattered signals. This was further improved in the “EnhanceOrSuppressFeatures” by enhancing the line structures of the network using the “Neurites” operation and “Tubeness” enhancement method. Images were thresholded using a two classes “Otsu” method. The thresholded images were converted into objects and mean fluorescence intensity (MFI) and occupied area were measured within these objects using the “MeasureImageIntensity” and “MeasureImageAreaOccupied” modules. Measured values were summarized as the mean across the whole image.

Furthermore, cell proliferation was evaluated from Ki-67 fluorescence signals. First, nuclei signals were enhanced in the “EnhanceOrSuppressFeatures” module by using the integrated feature type “Speckles”. DAPI-positive nuclei were identified in the “IdentifyPrimaryObjects” module and were thresholded by a two classes “Otsu” method. Subsequently, the Ki-67 image was masked in the “MaskImage” module. The previously identified nuclei objects served as a reference for the mask. The mean fluorescence intensity (MFI) of the Ki-67 mask was then measured within the nuclei objects.

### 4.12 Statistical analysis

Statistical analysis was performed using GraphPad Prism v10.4.1 (GraphPad Software, La Jolla, CA, USA). The applied statistical tests with multiple comparison are outlined in each figure legend. Significances are indicated as follows: *p < 0.05, **p < 0.01, ***p < 0.001, ****p < 0.0001. A p-value < 0.05 was considered statistically significant.

## Data availability statement

The data that support the findings of this study are available from the corresponding author upon reasonable request.

## Acknowledgments

A.S.M. was funded by the Deutsche Forschungsgemeinschaft (DFG, German Research Foundation) under Germany’s Excellence Strategy—EXC 2051—Project-ID 390713860.

A.S.M. was further supported by the DFG Collaborative Research Centre CRC 1278 “PolyTarget” (project ID316213987, subproject Z01, A05).

T.K. was funded by the Thüringer Aufbaubank (Germany) – 2021 SD0018 and 2023 SD0029.

M.A. was supported by a Scholarship from the Interdisciplinary Center of Clinical Research of the Jena University Hospital (Interdisziplinäres Zentrum für Klinische Forschung, IZKF Jena).

J.S. was supported by the Hermann Strauss Scholarship from the German Crohn’s Disease/Ulcerative Colitis Association (Deutsche Morbus Crohn/Colitis ulcerosa Vereinigung; DCCV e.V.).

The authors would like to thank Christian Becker for creating the rendered chip illustrations and intestinal model figures. In addition, we would like to acknowledge Sophie Besser, Daniela Schulz, and Tobias Vogt for their excellent technical lab support.

## Author contributions

T.K., J.S., K.G, M.R., A.S.M. contributed to conceptualization. T.K., M.A. contributed to methodology. T.K. contributed to investigation. T.K. contributed to formal analysis. T.K. contributed to visualization, T.K. contributed to writing - original draft preparation. T.K., M.A., J.S., K.G., M.R., A.S.M., contributed to writing - review and editing. K.G., M.R., A.S.M. contributed to supervision. K.G., A.S.M. contributed to project administration. M.R., A.S.M. contributed to funding acquisition.

