## Supplementary Material for "Secondary bile acid lithocholic acid ameliorates colitis-like inflammation in a human intestine-on-chip system"

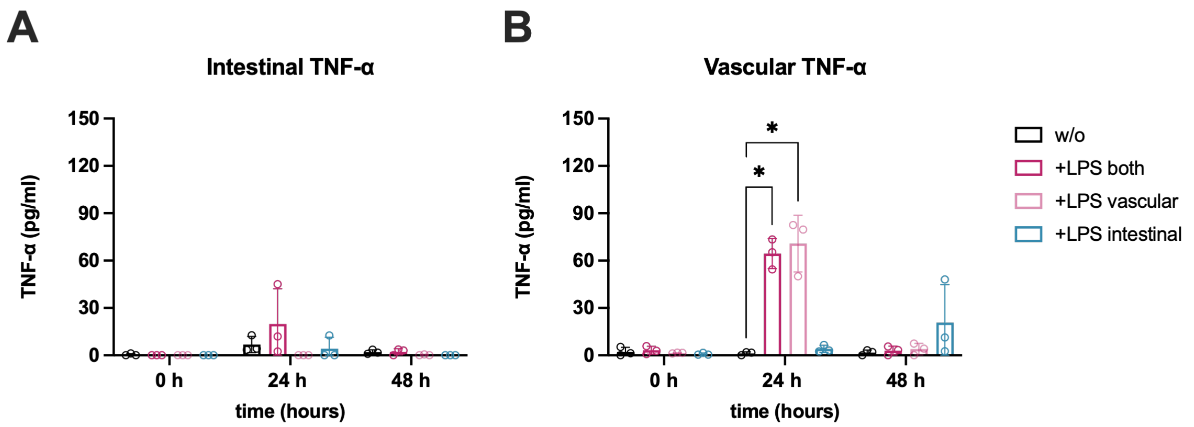


**Figure S1:** ***Pro-inflammatory cytokine response after LPS stimulation.*** *Measurement of TNF-α from intestinal* ***(A)*** *and vascular* ***(B)*** *medium supernatants in intestine-on-chip models treated without (-) or with (+) 100 ng/mL LPS in the indicated chip channels (both = vascular + intestinal channel, vascular = vascular channel, intestinal = intestinal channel) for up to 48 h. Bars represent mean ± SD of 3 independent experiments (n = 3) with 3 independent MDM donors. *p ≤ 0.05 (Two-way ANOVA with Dunnett's multiple comparisons test).*

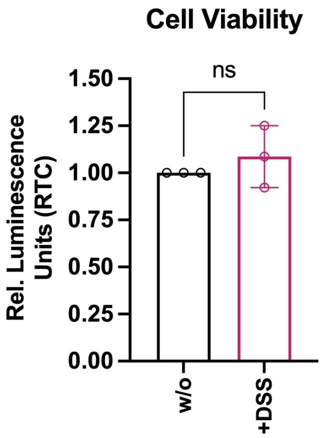


***Figure S2: DSS does not induce cytotoxicity in intestine-on-chip models.*** *Measurement of cell viability in untreated (w/o) or DSS-treated (+DSS) models after 48 h of treatment. Relative luminescence units are plotted as ratio to untreated control (RTC). Bars represent mean ± SD of 3 independent experiments (n = 3) with 3 independent MDM donors. ns = not significant (Paired two-tailed t test).*

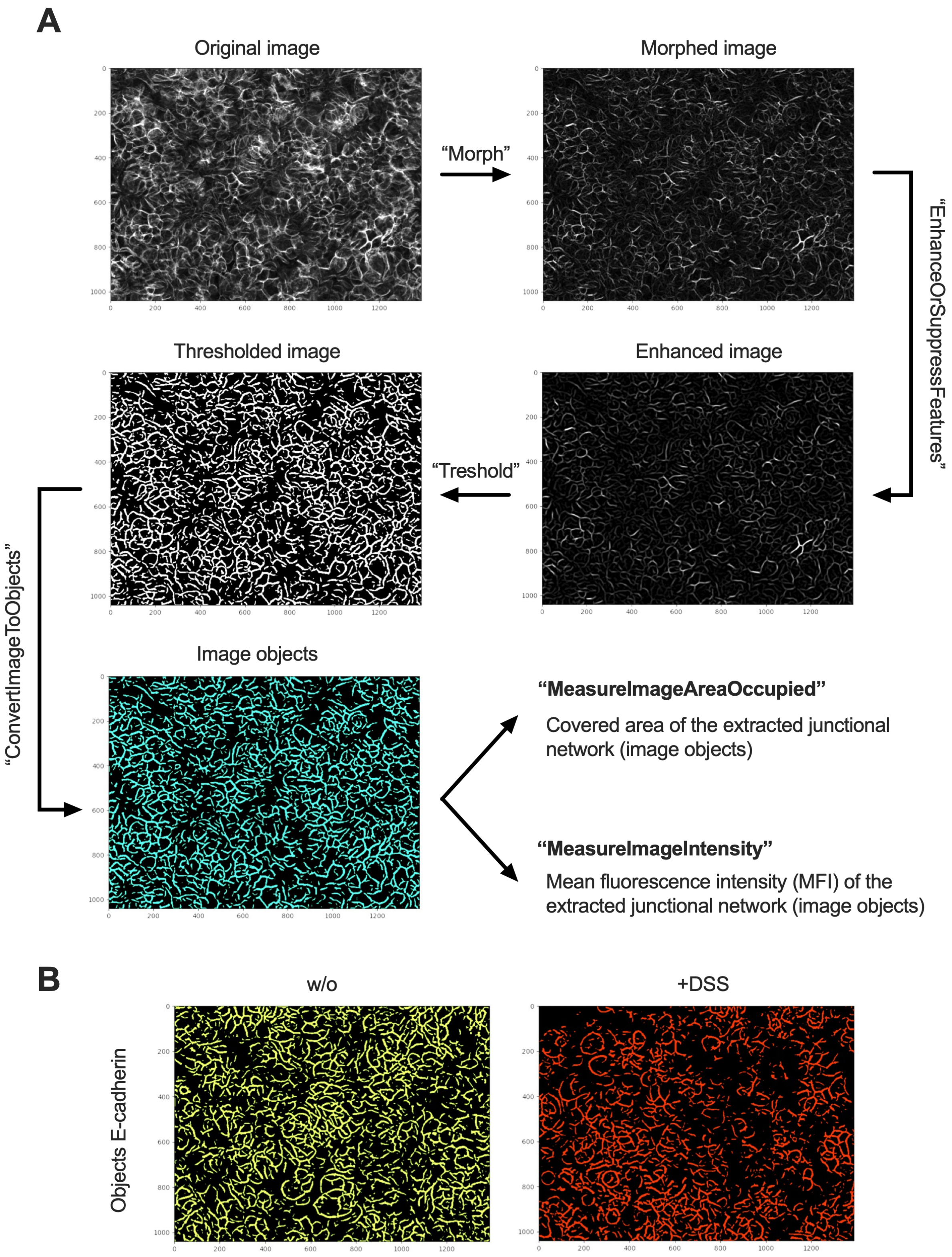


***Figure S3: Schematic illustration of the bioinformatic pipeline for quantification of junctional networks and adhesion molecules. (A)** Overview of the quantification pipeline with indicated CellProfiler modules. **(B)** Representative images of junctional network objects from untreated (w/o) and DSS-treated models.*

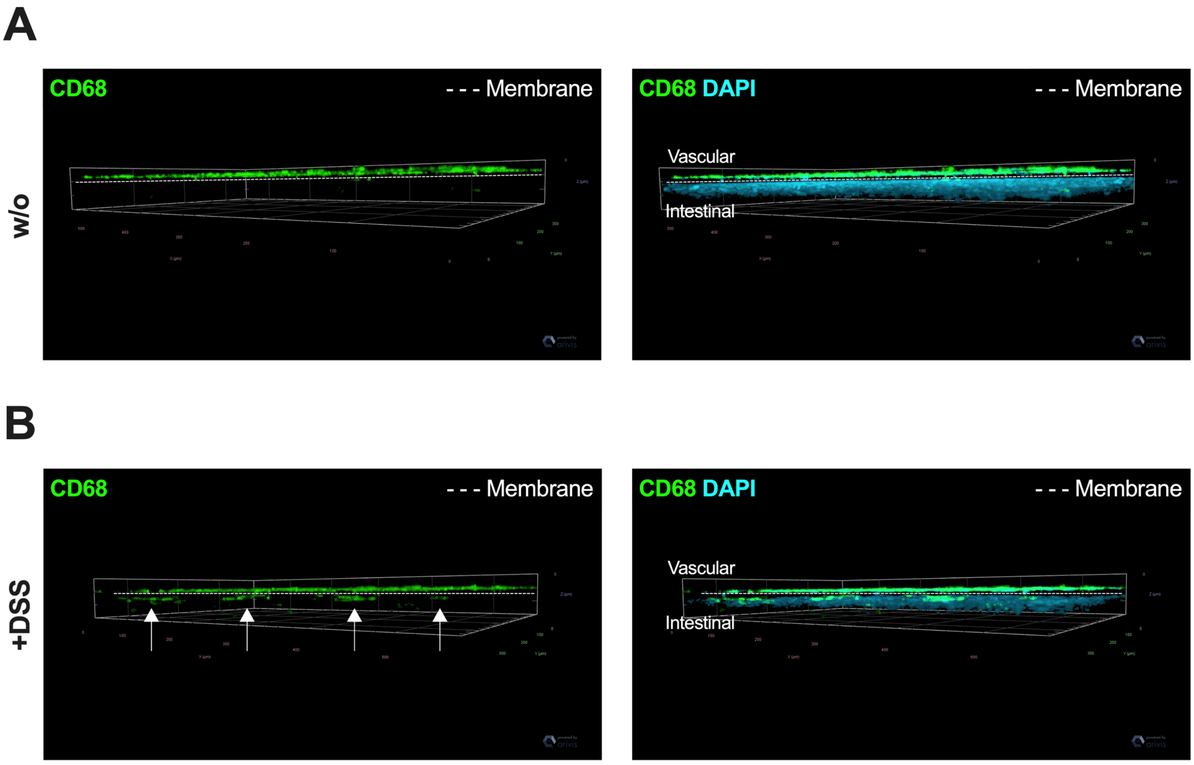


***Figure S4: MDMs infiltrate the intestinal tissue upon DSS-treatment.*** Representative 3D fluorescence reconstructions of **(A)** untreated intestine-on-chip models (w/o) or **(B)** DSS-treated models (+DSS) for 48 h. Macrophages were stained for CD68 (green). Vascular and intestinal cell nuclei were stained with DAPI (cyan). The integrated porous membrane is indicated with a dotted white line between the vascular and intestinal chip channel. White arrows indicate translocated CD68-positive MDMs **(B)**.

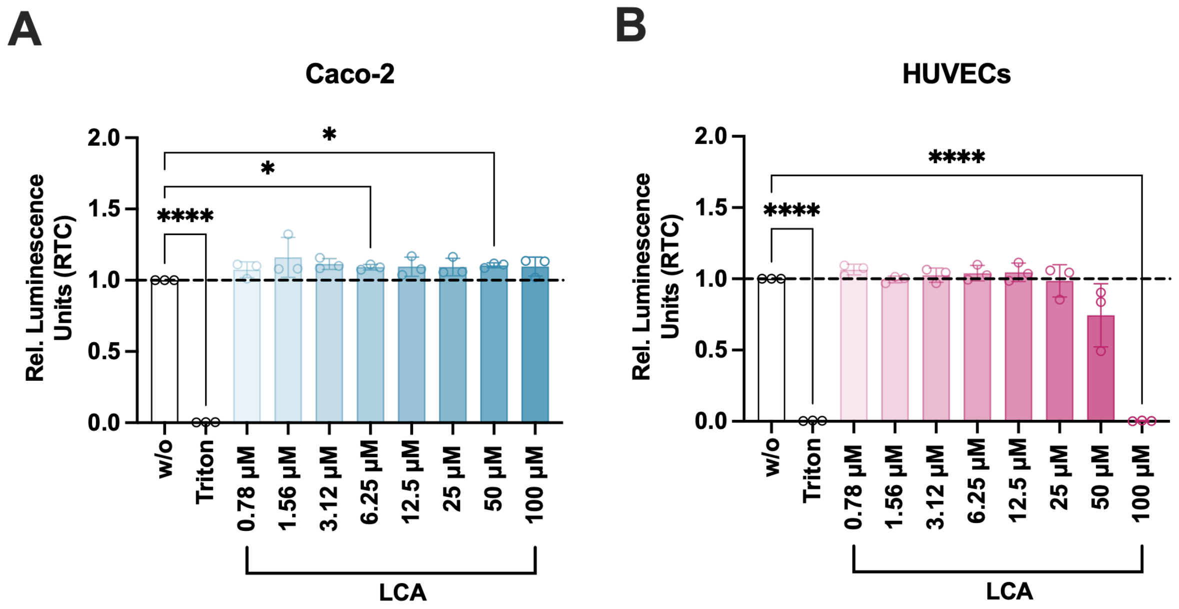


***Figure S5: Analysis of cell viability in HUVECs (A) and Caco-2 (B) monocultures after treatment with different concentrations of LCA for 72 h.*** *Relative luminescence units are plotted as ratio to untreated control (RTC). Bars represent mean ± SD of 3 independent experiments (n = 3). The dotted line represents the baseline of untreated cells (w/o). Triton X-100 (Triton) was used as positive control to reduce cell viability. *p ≤ 0.05, ****p ≤ 0.0001 (One-way ANOVA with Dunnett's multiple comparisons test).*

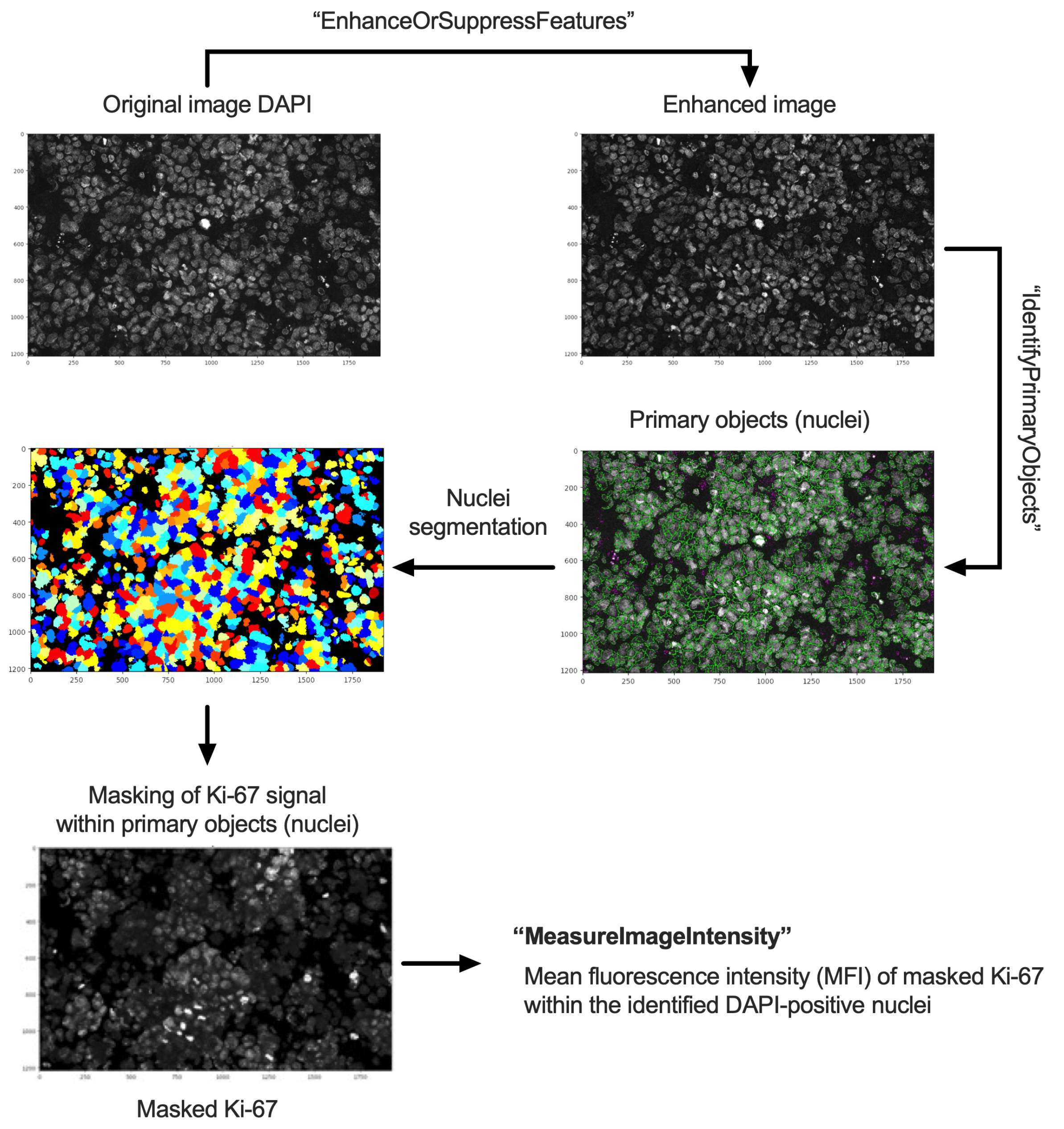


***Figure S6: Schematic representation of the bioinformatic pipeline for quantification of Ki-67 fluorescence signals.*** *The pipeline consists of different CellProfiler modules to identify DAPI-positive nuclei as primary objects and to subsequently measure the mean fluorescence intensity (MFI) of Ki-67 within the pre-identified nuclei.*

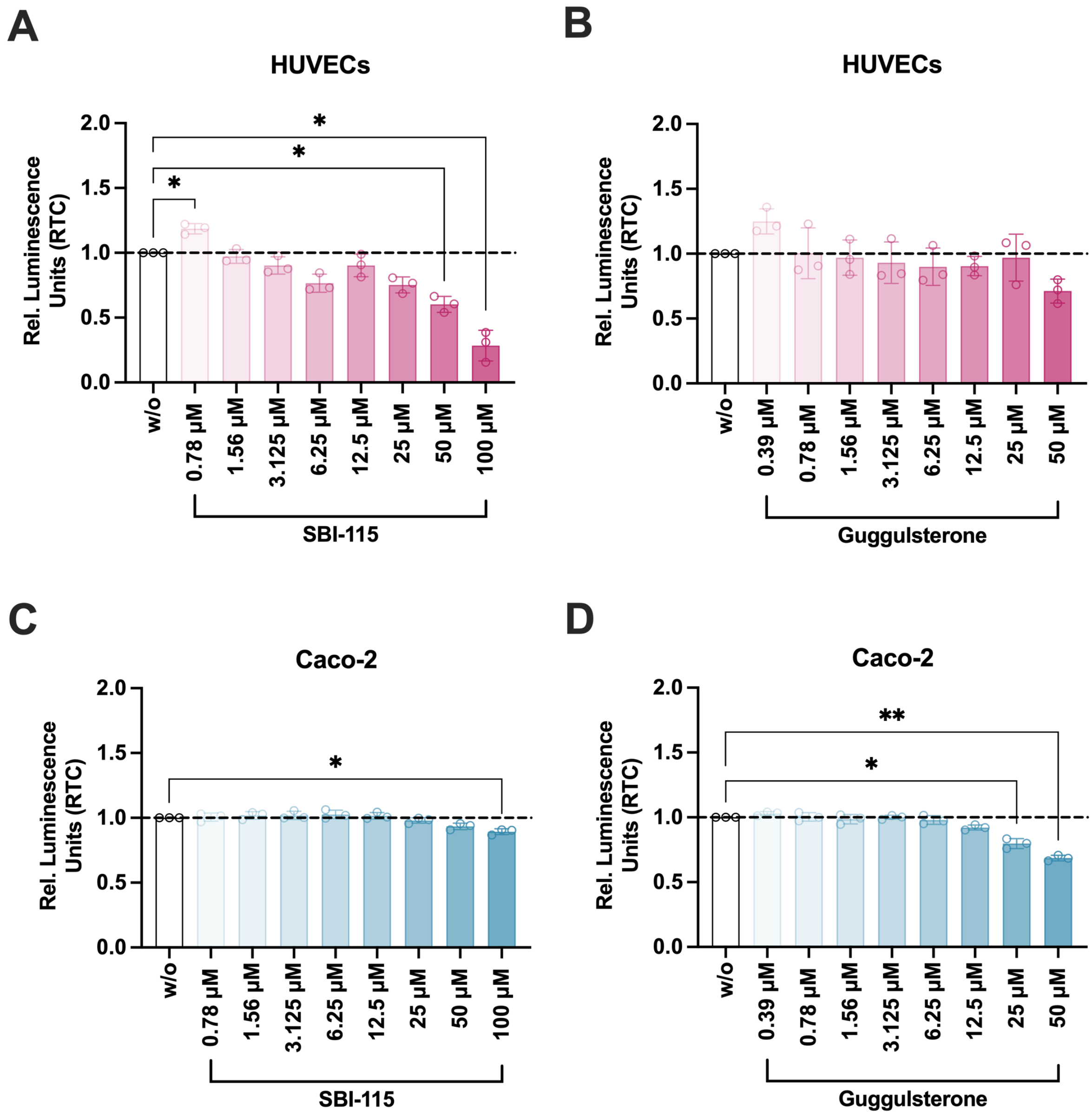


***Figure S7: Measurement of cell viability in monocultures treated with TGR5 and FXR antagonists.*** *Viability was measured in HUVECs **(A,B)** and Caco-2 **(C,D)** monocultures after treatment with SBI-115 **(A, C)** or guggulsterone **(B, D)** for 72 h. Relative luminescence units are plotted as ratio to untreated control (RTC). Bars represent mean ± SD of 3 independent experiments (n = 3). The dotted line represents the baseline of untreated cells (w/o). *p ≤ 0.05, **p ≤ 0.01 (One-way ANOVA with Dunnett's multiple comparisons test).*

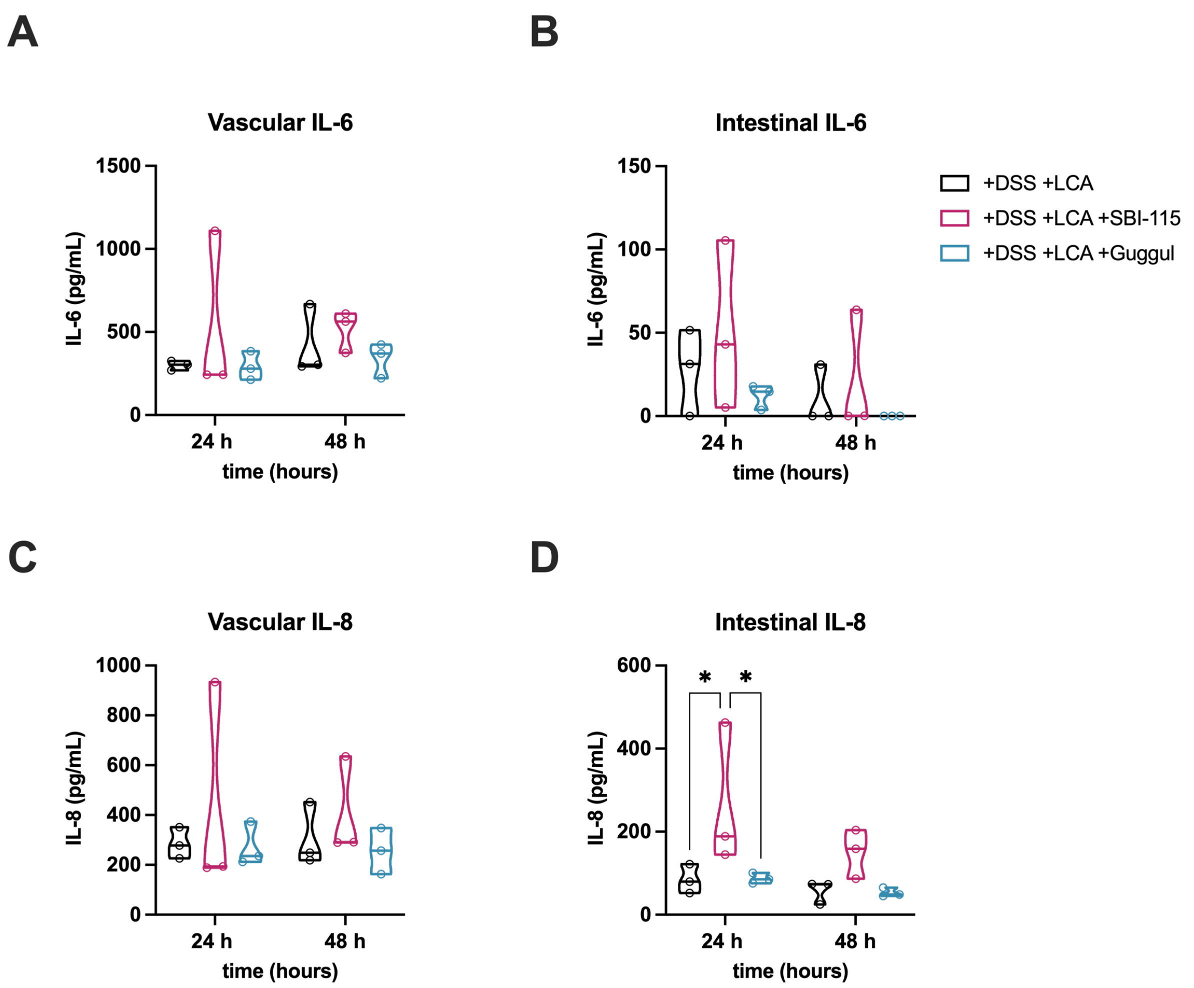


***Figure S8: Measurement of pro-inflammatory cytokine levels in response to TGR5 and FXR antagonism.*** *Analysis of pro-inflammatory IL-6 (**A-B**) and IL-8 (**C-D**) from vascular and intestinal medium supernatants of models treated with DSS/LCA (+DSS +LCA), DSS/LCA/SBI-115 (1 µM) (+DSS +LCA +SBI-115), and (+DSS +LCA +Guggul) DSS/LCA/guggulsterone (3 µM). Violin plots with indicated median and quartiles of 3 independent experiments (n = 3) with 3 individual MDM donors. *p ≤ 0.05 (Two-way ANOVA with Tukey's multiple comparisons test).*
